# Small animal brain surgery with neither a brain atlas nor a stereotaxic frame

**DOI:** 10.1101/2024.03.12.584491

**Authors:** Shaked Ron, Hadar Beeri, Ori Shinover, Noam M Tur, Jonathan Brokman, Ben Engelhard, Yoram Gutfreund

**Affiliations:** The Bruce and Ruth Rappaport Institue and Faculty of Medicine, the Technion, Haifa, Israel

## Abstract

Stereotaxic surgery is a cornerstone in brain research for the precise positioning of electrodes and probes, but its application is limited to species with available brain atlases and tailored stereotaxic frames. Addressing this limitation, we introduce an alternative technique for small animal brain surgery that requires neither an aligned brain atlas nor a standard stereotaxic frame. This method requires a high-contrast MRI scan of a specimen and access to a microCT scanner. The process involves attaching miniature markers to the skull, followed by CT scanning of the head. Subsequently, MRI and CT images are co-registered using standard image processing software and the targets for recordings in the brain are marked in the MRI image. During surgery, the animal’s head is stabilized in any convenient orientation, and the probe’s 3D position and angle are tracked using a multi-camera system. We have developed a software that utilizes the on-skull markers as fiducial points to align the CT/MRI 3D model with the surgical positioning system, and in turn instructs the surgeon how to move the probe to reach the targets within the brain. Our technique allows the execution of insertion tracks connecting two points in the brain. We successfully applied this method for neuropixels probe positioning in owls, quails, and mice, demonstrating its versatility and its potential to open new avenues for research in non-standard and novel animal models.

## Introduction

Systems neuroscience often employs stereotaxic surgeries for positioning electrodes, probes, pipettes, and optical fibers within the brain. This technique relies on two essential factors: standard head positioning that is reproducible across animals and labs and a species-specific brain atlas aligned with the standard positioning (De Vloo & Nuttin, 2019). The coordinates from established landmarks are extracted from the atlas (Paxinos *et al*., 1985) to guide instruments to targeted brain areas. Over the years, advancements like digital 3D atlases, sophisticated stereotaxic frames, and motorized control systems have enhanced precision and reliability (Löffler *et al*., 2009; Wang *et al*., 2020; Kleven *et al*., 2023). However, the major drawback of the technique is that it can be performed only in animal species for which stereotaxic tools have been developed and are available. This restricts the availability of the technique to a hand-full of common laboratory animal models in neuroscience such as the mouse, rat, zebra-finch, pigeon etc. While there is a growing interest in expanding animal models in neuroscience to include unique, behaviorally interesting species (Keifer & Summers, 2016), the research into non-standard species is hindered by the lack of matched stereotaxic tools.

A potential solution is to produce a brain atlas and stereotaxic techniques specific for each animal model (Karoubi *et al*., 2016; Eilam-Altstadter *et al*., 2021). However, the resources and years of development required are usually beyond the capacity of most research laboratories. To address this challenge, we have developed an alternative technique that eliminates the need for standard head holding positioning devices or aligned brain atlases. This method, instead, relies on the capability to perform a CT scan on a live animal. While this is a limitation, ‘small animals’ micro CT capabilities are increasingly standard in research facilities, offering a non-invasive and cost-effective solution (Clark & Badea, 2021). Notably, CT scanners are suitable for a wide range of animal species, making this technique applicable to various species without existing brain atlases, including species never previously used in neuroscience research.

This technique is based on miniature markers affixed to the skull, serving as fiducial markers. They enable the transformation of the brain’s coordinate system from a CT/MRI 3D model to that of the surgical probe positioning system. We have developed software that calculates and utilizes this transformation, instructing the surgeon how to guide the probe’s movements. Drawing inspiration from human brain surgery methods (Zhou *et al*., 2017), we have integrated a 3D motion tracking device. This device precisely measures the surgical probe’s angle and XYZ position. The gathered data is then used to calculate the deviation of the probe from the target, allowing for real-time adjustments in both angle and position.

In summary, we present a solution to the challenge of stereotaxic surgery in small animals lacking an established brain atlas. Our system allows the execution of planned insertion tracks connecting two points within the brain. Given its relatively low cost and the increasing availability of the necessary facilities, this technique could serve as an alternative or complement to standard stereotaxic methods in mice and other well-established animal models. In this paper, we comprehensively detail our method and demonstrate its reliability. We highlight its successful application in guiding the insertion of neuropixels probes (Jun *et al*., 2017) for acute recordings in the entopallium of barn owls, as well as in the chronic implantation of probes in the thalamus of quails and the Ventral Tegmental Area (VTA) of mice.

## Methods

### Animals

Two adult barn owls, two quails and two mice were used in this study. Animals were born and raised in captivity and housed in aviaries and cages in accordance with the NIH guide for the care and use of laboratory animals. All procedures were approved by the Technion’s Institutional Animal Care and Use Committee. All surgical procedures were performed under full anesthesia. No painful procedures were carried out during the recording sessions.

### Ex-vivo MRI scan

One adult female quail (*Coturnix japonica*) and one adult male barn-owl (*Tyto alba*) were used for high resolution structural MRI scan, following a protocol modified from (Güntürkün *et al*., 2013). The animals were euthanized (Heparin and Pentobarbital) and perfused with a standard PBS solution (0.9%, Sigma-Aldrich, St. Louis, MO, USA), followed by PFA (4%, Sigma-Aldrich) mixed with gadoteric acid (Cyclolux 0.5 mmol/ml, Sanochemia Pharmazeutica AG, Austria) in a 1% solution. Immediately after perfusion, animals were decapitated and heads were put in the same solution for at least 5 days in 4°C. Prior to scanning, heads were placed inside a plastic tube filled with a magnetically inert fluid (Fomblin, YL VAC 06/6, Solvay, Italy). Special care was given to remove all air bubbles from the sealed tube.

Samples were inserted to a 9.4T horizontal MRI system (Bruker Biospec, Ettlingen, Germany) interface with Avance III console, using a cylindrical volume coil (86 mm inner diameter) for RF transmission and a surface coil (20 mm diameter) for detection. 3D T_2_-weighted images were acquired using a Rapid Acquisition with Relaxation Enhancement sequence (RARE), at 100 um isotropic resolution, repletion time (TR) = 350 ms, Echo time (TE) = 17.2 ms and RARE factor = 4. Field of view (FOV) = 25.6 x 25.6 x 18 mm^3^, matrix size = 256 x 256 x 180, number of averages = 4, and total scan time of 4.5 hours.

### Surgical and CT scanning procedures

The surgical procedure is divided into two stages (Figure 1 illustrates the workflow of the procedure). The aim of the first surgery is to attach at least four fiducial markers to the skull that can be observed in the CT and in addition can be observed and measured during the surgery. It is possible and desired to use the first surgery to implant head posts, ground screws etc., to save valuable time in the second surgery. However, having relatively large metal bodies will contaminate the CT image and may induce inaccuracies in the MRI/CT alignment procedure and/or the identification of the markers. In barn owls we use plastic head-posts and therefore we implant these in the first surgery. In the quails and mice, because of their small head we prefer to insert the ground screws after the CT, in the second surgery.

**Figure 1.**
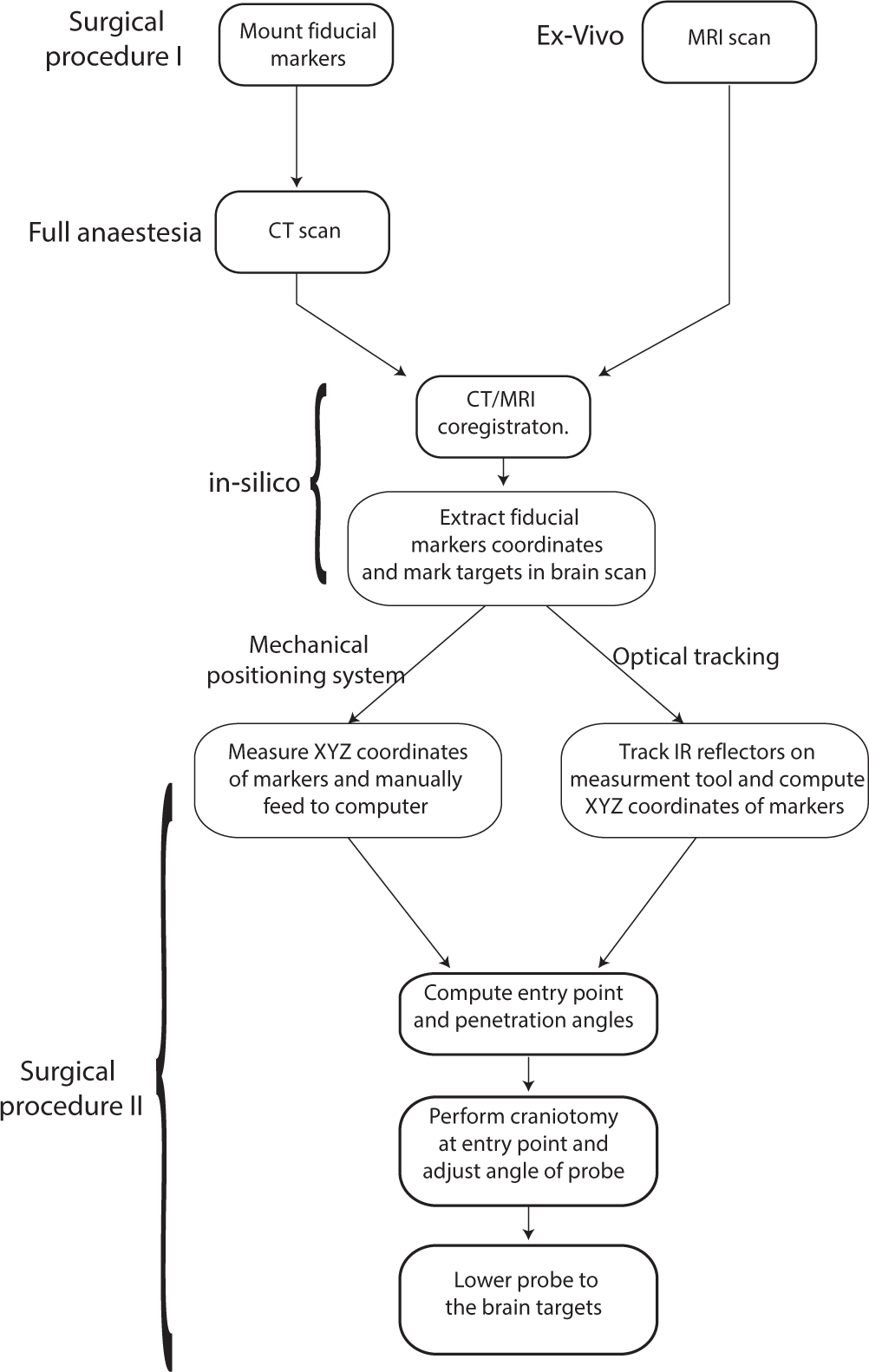
Flow chart of the procedure. The method requires three stages: An initial stage to attach the markers and to perform a CT scan of the head. A second stage which is performed on the computer, to align the MRI with the CT 3D volume. The third stage is the surgical procedure to conduct a craniotomy and lower the electrode onto its brain targets. This stage is guided by the software which uses the fiducial markers from the CT to compute the penetration track (position and angle).

#### Procedure of first surgery

The birds (owls and quails) and mice are anesthetized with isoflurane using standard protocols for each species. The animals are positioned on a heat blanket and lidocaine (Lidocaine HCl 2% and Epinephrine) is injected locally at the incision site. Then an incision of the skin is made to expose the skull bone. The exposed skull is scrubbed clean and dried and a thin layer of metabond (C&B Metabond adhesive) is spread over the exposed skull. Veterinary tissue glue (Vetbond) is applied to the edges of the cut skin. Then the markers are glued in place. We used two different approaches for fiducial markers and both worked equally well: 1) Metal markers - a tungsten wire 150 microns thick is cut with a fine cutter to four pieces of about 1 mm each. We put four drops of dental cement at different places on the exposed metabond layer and, before curing, the wires are imbedded in the cement in a vertical orientation. The markers need to form a unique rigid body with 6 DOF. The markers should not be on a single plane and ideally, spread as much as possible. We make sure that the markers are not on a single plane by building different heights with dental cement. After the cement is fully cured, under the binocular, a fine driller is used to expose the dorsal tip of each marker.

(2) Plastic frame - We designed a 3D-printed plastic rectangular frame with four cone-shape holes of 1 mm diameter and 1 mm depth. The holes are spread along the edges of the rectangular at different heights to form a unique rigid-body (Fig. 2A). The frame is glued to the metabond layer with the cone shape holes pointing upwards. The holes in the plastic are visible in the CT image and can provide convenient landmarks during the surgery. However, for a higher contrast of the fiducial markers in the CT, it is recommended to insert metal wires in the holes and glue with epoxy or dental cement (Fig. 2B). After the surgery, the animal recovers for several hours in a heated chamber and is returned to its home cage.

**Figure 2.**
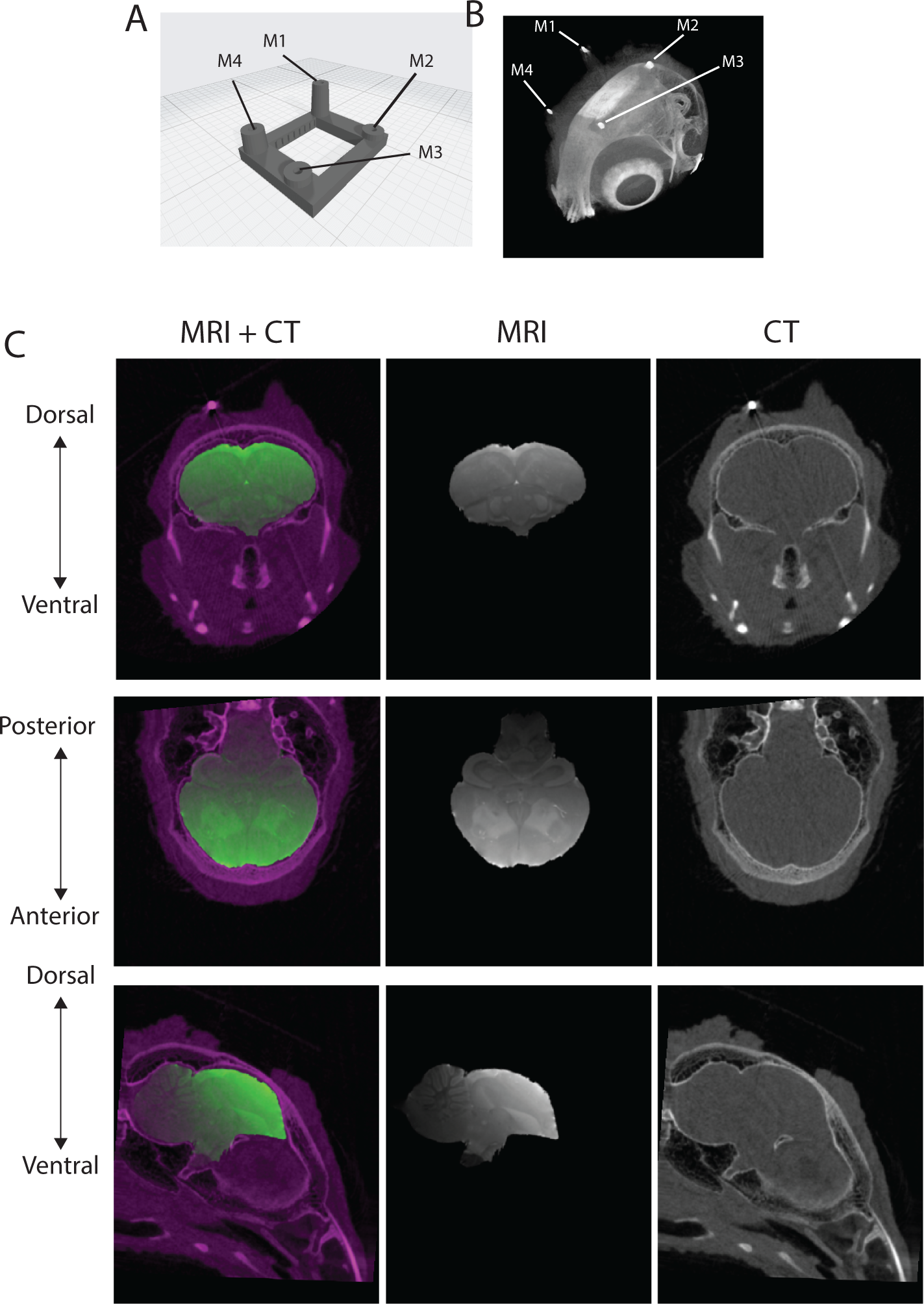
MRI and CT co-registration. A. The design of a 3D printed frame that is glued to the skull in the first surgical procedure. Metal wires are inserted into the four cone shaped holes to serve as fiducial markers. B. A 3D CT image showing the upper part of the quail’s skull. The four metal markers in the printed frame are clearly visible (M1 to M4). C. MRI volume of the quail’s brain is transformed to align with the quail’s skull in the CT volume, as described in Methods. The first column shows aligned MRI sections, in a coronal, sagital and transverse planes (green), superimposed on the coresponding CT images (purple). The second column shows the corresponding MRI images alone. The third column shows the corresponding CT images alone.

#### CT scan

About a week after the first surgery, the animals are anesthetized with intramuscular injection of ketamine hydrochloride (10%) and xylazine (2%), in owls 20 mg/kg and 4 mg/kg respectively, in quails 70 mg/Kg and 14mg/Kg respectively, in mice 80 mg/Kg and 8mg/Kg respectively. After anesthesia, the animals are restrained inside an X-ray tube for small animal (diameter ∼7 cm). Head pose in the tube is fixed by inserting pieces of X-ray inert-foam around the beak and securing with medical tape. CT scans are conducted in a Bruker micro CT scanner for animal research, using the following parameters:(a) MIS-BIC-2kVe configuration, (b) Al-Cu filter, (c) exposure time 240ms, (d) 80kV, 58uA, (e) 84µm voxel resolution (isometric volume), Rotation by 1.0 degree for 360 degrees, (g) Frame resolution of 504X306 pixels. Total time of scan is about 6 minutes. 3D volumes are reconstructed using a standard software – NRecon (version 1.7.4.6, Bruker) and then processed in common image analysis software packages (MIPAV & ImageJ, NIH).

#### Alignment of CT and MRI model

The MRI 3D image has to be scaled and aligned to fit the CT scan (volume registration (Li *et al*., 2008)). Numerous image processing tools offer volume registration (Lee *et al*., 2015; Fernandez & Moisy, 2020). We implemented the registration processes with an open source software package called Fijiyama (https://imagej.net/plugins/fijiyama) or with matlab *imregtform* funcrion. Fijiyama is a generic tool for 3D image series registration which is commonly used for X-ray MRI registrations. The two datasets are preprocessed and save as 3D TIF format images. These 3D images are loaded to Fijiyama. The alignment process is a two-stage process: a manual and an automatic stage. First, software tools are designed to allow manual registration, bringing the two datasets as close as possible by manually changing scales, orientations and positions. This first stage results with a rough superposition of the two data sets which serves as the initial condition for the next automatic step. In the next step, build-in algorithms find the optimal registration of the MRI image with the reference CT image. Finally, we perform a manual inspection of the superimposed images to ensure that the MRI is satisfactorily aligned with the CT (Figure 2C). After the registration is finished, the XYZ coordinates of the four fiducial markers (dorsal edges of metal wires or center of coin-shaped holes in plastic frame) are identified in the CT (Figure 2B) and their coordinates are inserted in the Matlab code. The brain targets are identified in the registered MRI images and also inserted in the code (see supplementary appendix for code and guidelines).

#### Second surgical procedure

The aim of the second surgery is to position the electrodes in the brain at the correct location and angle. The animals are anesthetized with isoflurane as described above, then the head is fixed to maintain a stable position throughout the surgery. However, the position and orientation of the head is not critical and can be maintained at any convenient orientation. For owls we hold the head in a fix and repeatable position via the head post that is attached to the skull in the first surgery. In the quails we use ear bars (rat ear bars) and a custom beak holder to fix the head. At this stage, depending on the available hardware, there are two options for carrying out the surgical procedure: one is to use a mechanical XYZ positioning system, the second is to use a 3D video-based tracking system. It is also possible to combine both options. For example, use an XYZ positioning system to measure the coordinates of the fiducial markers and a 3D tracking system to adjust online the angle of the probe. Below we describe both options to give the user maximal flexibility.

Mechanical XYZ positioning system: Any XYZ manipulator with a position readout can be used. We use Stoelting 3 arm digital manipulator. To measure the XYZ coordinates of the fiducial markers a syringe needle (25 G) is attached to the vertical arm of the manipulator. Under binocular inspection the tip of the needle is moved to touch the first of the markers. The readout of the XYZ positioning system is zeroed and this marker becomes the origin of the axes (0,0,0 point). Then the needle is moved to touch the second marker and the XYZ distances from the first point are measured. This is repeated for the remaining two points and the measurements are fed to the computer. We now have the positions of the fiducial markers in the coordinate system of the surgical environment, which are used to calculate the estimated positions of the desired brain targets in the surgical environment (see below *MatLab software and computations*). The continuation of the procedure depends on the type of surgery: single brain target (vertical penetration) or two brain targets (angled penetration). In the single target surgery, the XYZ position of the brain target is provided by the software, the needle is moved to the correct XY position and carefully lowered until touching the exposed metabond surface to mark the center of the craniotomy. A craniotomy is performed to expose the brain surface and the probe is then inserted vertically at the correct XY coordinates until reaching the desired depth.

For two brain targets, the software provides the estimated XYZ position of the entry point on the skull and the desired angle of the probe (see below *<u>MatLab software and computations</u>* program details). The needle is then moved to the predicted entry point to mark the position of craniotomy. The craniotomy is performed to expose the brain surface. Now the probe is attached to the XYZ positioning system and the angle of penetration is adjusted to fit the desired angle. Note that most available XYZ manipulators do not have an option to adjust a precise 3D angle. The Stoelting arm that we are using, for example, is limited to control only the yaw or pitch angle (but not both) and at a precision of about 2 degrees. The use of a ball joint and 3D video tracking solves this limitation.

#### 3D video tracking

3D optical tracking of IR reflectors is a common solution for motion tracking, used in medical devices, robotics and animal research (Dubois & Bresciani, 2018; Singh *et al*., 2022). There are several commercial devices but it is also possible to customize a home-made tracking device (Yang *et al*., 2012). We use a commercial motion tracker (OptiTrack; https://optitrack.com/). Our system is composed of 3-4 high-quality infrared (IR) sensitive cameras (Prime^x^13), a laptop and the Motive software (OptiTrack). Cameras are scattered in the surgery room (Figure 3A) and connected to a laptop through an ethernet network switch. After calibration of the cameras, the positioning accuracy is estimated by the Motive software to about ± 0.1 mm. To achieve maximal maneuverability of the probe to be inserted, we use an articulated arm with two ball joints, locked with a single screw (Articulated holder, NOGA Engineering, Israel). For additional fine movements of the probe, an XYZ micromanipulator with an oil hydraulic control of motion along the z axis (MN-153, Narishige Group, Japan) is attached to the NOGA arm (Figure 3B). We devised a 3D printed measurement tool (with three spherical IR reflectors of 6.3 mm diameter and a 25G needle at the tip. The reflectors form a straight line with known distances to the tip of the needle (Figure 3C). The measurement tool is attached to the manipulator (Figure 3B) and its tip is moved, under binocular inspection, to touch the fiducial markers one by one. For each marker the MatLab program reads the XYZ positions of the IR reflectors from the Motive software and computes the XYZ coordinates of the fiducial markers. Then the program finds the rigid body transformation that best transforms the CT based model to the surgical coordinate system and use the transformation matrix to estimate the coordinates of the brain targets, the craniotomy location and the angle of penetration (see below *MatLab software and computations)*. The craniotomy is then performed to expose the brain surface. The angles of the measurement tools are adjusted to fit the desired angles in the posterior-anterior plane and the lateral medial plane. For an accurate and easy adjustment, the program continuously reads the positions of the IR reflectors, computes and displays in real time the current angle and position relative to the desired angle and position. After the correct angle is achieved, the measurement tool is replaced with the probe holder. The probe is then driven with the fine oil hydraulic micromanipulator to penetrate the brain at the center of the craniotomy, all the way to hit the deep brain target.

**Figure 3.**
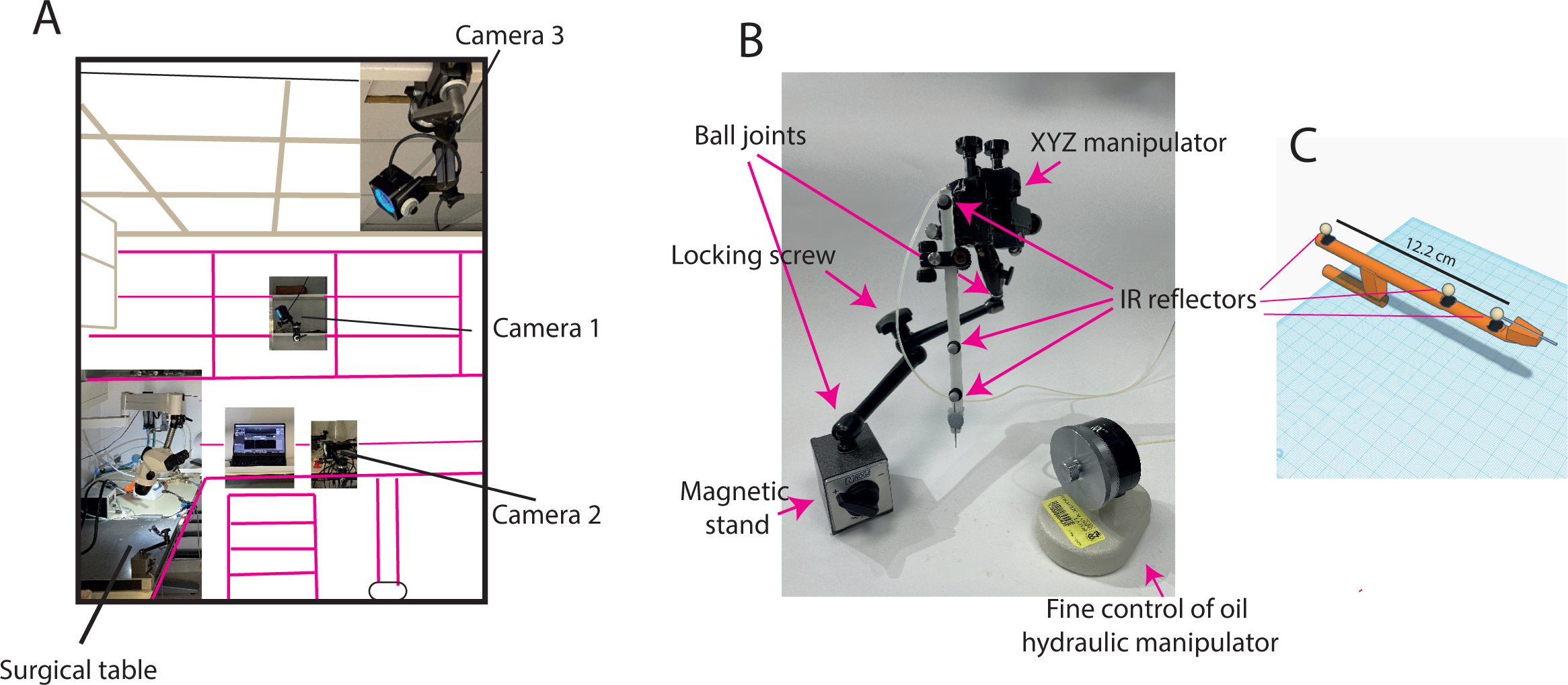
The surgical setup. A. A scheme of the surgical room showing the positioning of three video cameras for optical tracking. Each camera is equipped with a ring of IR leds to illuminate the IR reflectors on the measurement tool. The cameras are connected to the laptop with ethernet cables. B. The XYZ manipulator attached to the end of a NOGA arm allows a high flexibility of positioning the probe at any desired angle in 3D. Movement along the probe axis is controlled by a fine oil hydraulic manipulator. C. A sketch of the measurtement tool. Three spherical IR reflectors with known distances from the tip are used to define the position and angle of the probe.

### MatLab software and computations

We developed a MatLab program to accompany the surgical procedure (see Supplementary Appendix for code and user guide). Before the surgery, the coordinates of the fiducial markers in the CT, the two brain targets and the point of entry need to be inserted in the code. There are good, freely available tools for viewing and navigating 3D images. Therefore, we did not develop our own tool for marking the CT and/or MRI images. Instead, we use MIPAV (https://mipav.cit.nih.gov/) to extract the coordinates in voxels of the metal markers, in the CT, and the brain targets, in the registered MRI. After the two targets in the brain are marked, the entry point on the skull can be estimated. For this, we developed a MatLab tool that displays the line connecting the two brain targets on the CT image (Figure 4). The user can select the intersection point of the penetration line with the surface of the skull (Figure 4) to mark the coordinates of the entry point, the position on the skull for performing a craniotomy. If only one brain target is selected, the penetration is vertical and there is no need to mark an entry point.

**Figure 4.**
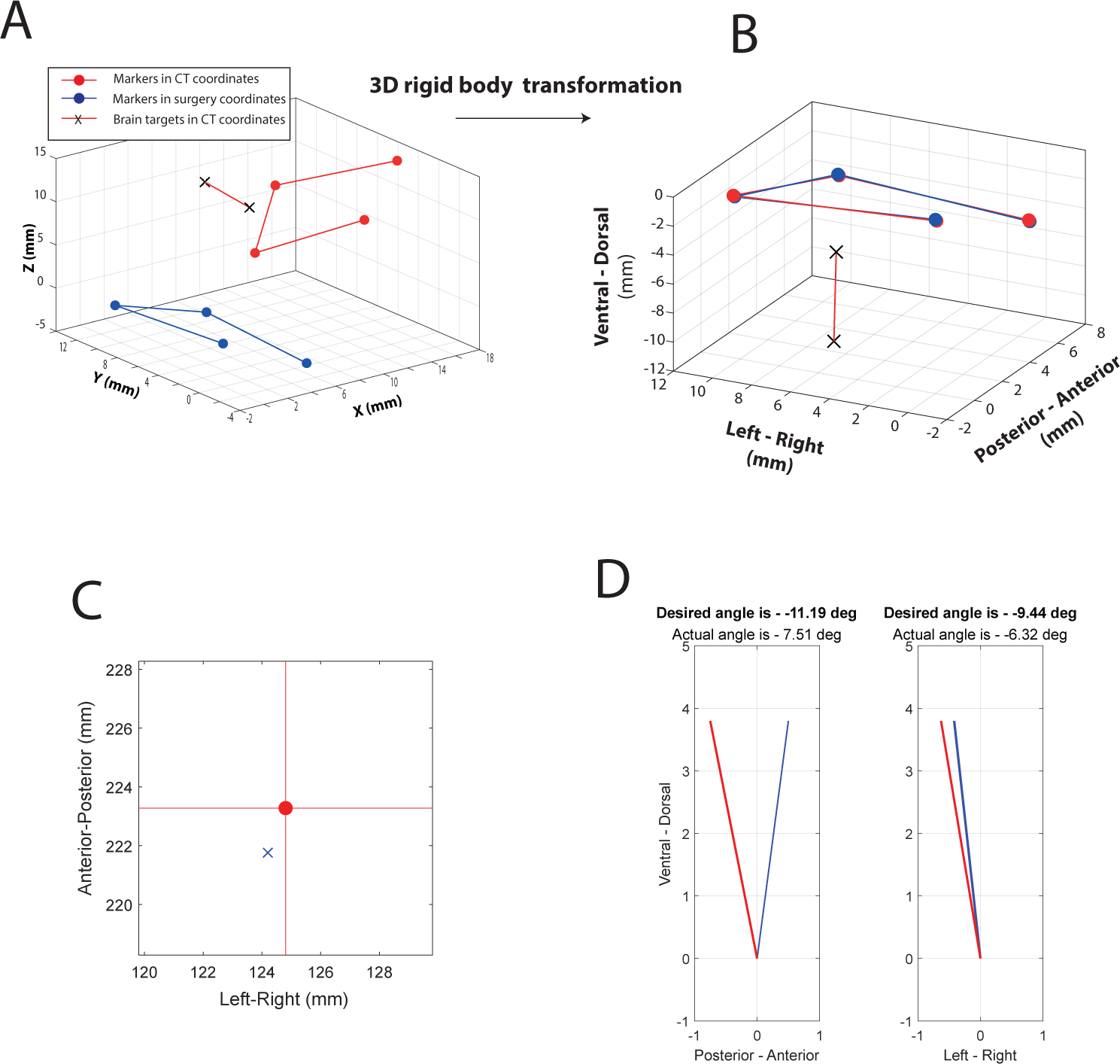
Graphs displayed by the software to guide the surgery. A. A 3D plot showing the position of the four fiducial markers in surgery coordinates (blue dots). On the same reference frame the markers (red dots) and two brain targets (black Xs) are ploted in the CT coordinates. B. The CT coordinate system is transformed, as described in Methods, to optimally fit the CT position of the markers with the real position of the markers in the surgery. The display in B superimposes the transformed CT markers over the real positions of the markers and shows the prediction of the targets position in space. This display is used to assess the quality of the fit before moving to the next stage. C. The red dot at the center of the graph is the XY position of the calculated point of entry on the skull. The blue x displays the position of the tip of the probe, which is updated in real time. By gently moving the probe the surgeon aims to bring the blue X to the red dot. D. The angles of penetration in the posterior-anterior plane and in the left-right plane are calculated and displayed as red lines oriented at the calculated angles, to mimick the desired positioning of the probe in real-space. The blue lines display the current orientation of the probe on the corresponding planes. This orientation of the blue lines is updated in real time and the aim of the surgeon is to align the blue lines with the red lines before penetrating the brain.

Towards the surgery, the user can choose to use optical tracking or not. If choosing not to use optical tracking, the coordinates of the fiducial markers are measured as described above using the mechanical XYZ positioning system and entered manually to the program. In this case the program is generic and can be used with any holding manipulator. On the other hand, if choosing optical tracking mode, the program calls NatNet SDK library for streaming data from the Motive software (https://optitrack.com/software/natnet-sdk/). If using a different tracking system, the code needs to be modified accordingly.

In the optical tracking mode, the program instructs the user to position the tip of the measurement tool on the first fiducial marker, checks that the three IR reflectors are identified and computes the XYZ coordinates of the fiducial marker (V_fm_):

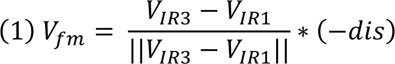

Where *V_IR1_* and *V_IR3_* are the 3D vectors of the lower and upper IR reflectors respectively, and *dis* is the Euclidian distance between the center of the lower IR reflector and the tip of the needle. For better positioning of markers, the program averages 100 measurements of each marker. This procedure is repeated for all four markers. At the end of this stage the program holds a variable *m* (4 x 3 matrix) with the coordinates of the four markers in the CT coordinate system in voxels numbers (extracted from the CT image), and a variable *p* (4 x 3 matrix) with the coordinates of the four markers in the surgery coordinate system in mm (obtained from the optitrack system). The CT coordinates are multiplied by the scale factor (k) to bring to a common unit (Fig. 4A);

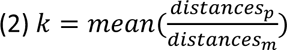

where distances_p_ and distances_m_ are the Euclidian distances between all points in *p* and in *m* respectively.

Next, the program finds the best transformation, represented by rotation matrix and translation vector (R, T respectively), between the two coordinate systems that align the four markers together (rigid-body transformation problem). We adopted an algorithm introduced by Arun et al., (Arun *et al*., 1987) and evaluated by (Eggert *et al*., 1997) which applies a singular value decomposition (SVD) to compute R and T that minimize the error term:

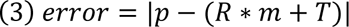

At this stage, the program displays a 3D plot showing the transformed markers, brain targets and point of entry in the coordinate system of the surgery (Figure 4B). This display is important for visually assessing the quality of the fit. The program also provides the coordinates of the entry point and the two penetration angles (angle on the, sagittal plane (XZ plane) and angle on the coronal plane (YZ plane)):

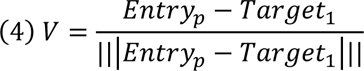

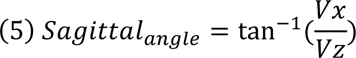

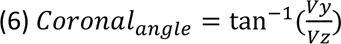

Where *Entry_p_* is the coordinates of the entry point, *Target_1_* is the coordinates of the deeper of the two brain targets. V is the unit vector from Target_1_ pointing to *Entry_p_*. Vx, Vy and Vz are the x,y and z components of V.

If the quality of the fit is satisfying, the user can move to the next stage where the position of the tip of the probe is calculated and displayed in real time (Figure 4C). Additionally, the angles of the probe are calculated and displayed in real-time (Figure 4D). The tip of the probe is calculated as in equation 1. The angles (sagittal and coronal) are calculated as in equations 4-6 but the entry point and target point are replaced by the position of the upper IR reflector and the position of the lower IR reflector on the probe, respectively.

### Histloogy

Immediately before penetrating the brain, neuropixels probes were dipped in DiI solution (Invitrogen). After the experiments, the animals were perfused transcardially with PBS followed by 4% paraformaldehyde. The brains were removed, maintained in paraformaldehyde+sucrose solution for a week and sliced with a cryostat (50 μm coronal slices). Slices were mounted, stained with DAPI (invirtogen) and imaged (slide scanner, NIKON) using DAPI and DiI filters.

## Results

### *Error assessmen -* Statistical simulation

To quantitatively assess the method’s accuracy, we simulated a noisy sample of four fiducial markers. This involved drawing marker positions from a normal distribution centered around the original markers’ positions on the 3D printed frame. We selected an isotropic standard deviation of 100 µm for the normal distribution because this value represents the estimated error in positioning a single marker using our OptiTrack setup. This error is also comparable to the voxel size in the CT image, which is 84 µm. We generated two sets of simulated imperfect marker positions: one to mimic the positioning error of the fiducial markers in the CT and another to mimic the positioning error of the fiducial markers in surgery (accuracy of optical or mechanical positioning).

Using these two sets of points, we apply the algorithm described above to calculate the transformation and rotation matrix that optimally aligns one set of simulated points to the other. The transformation is then applied to a target point, and the estimated error is calculated as the Euclidean distance between the transformed target and the original target.

With this approach, we simulated 2D grids of 41×41 target points (with a 250 µm interval between targets) on five parallel planes, 2, 4, 6, 8 and 10 mm below the plane of the frame. For each virtual target, we computed the transformation error as outlined above. This process was repeated 100 times and averaged to create accuracy maps (Fig. 5B). The simulation revealed that positioning accuracy decreases with distance from the frame’s center, both along the Z axis and in the X,Y plane. This decrease is expected as orientation errors typically amplify with distance from the rigid body. The maximal expected error, located 5 mm to the side and 10 mm below the frame’s center, was about 350 µm. The expected error for a penetration of 10 mm directly below the frame’s center (equivalent to the full length of a Neuropixel probe) was about 290 µm (Figure 5B).

**Figure 5.**
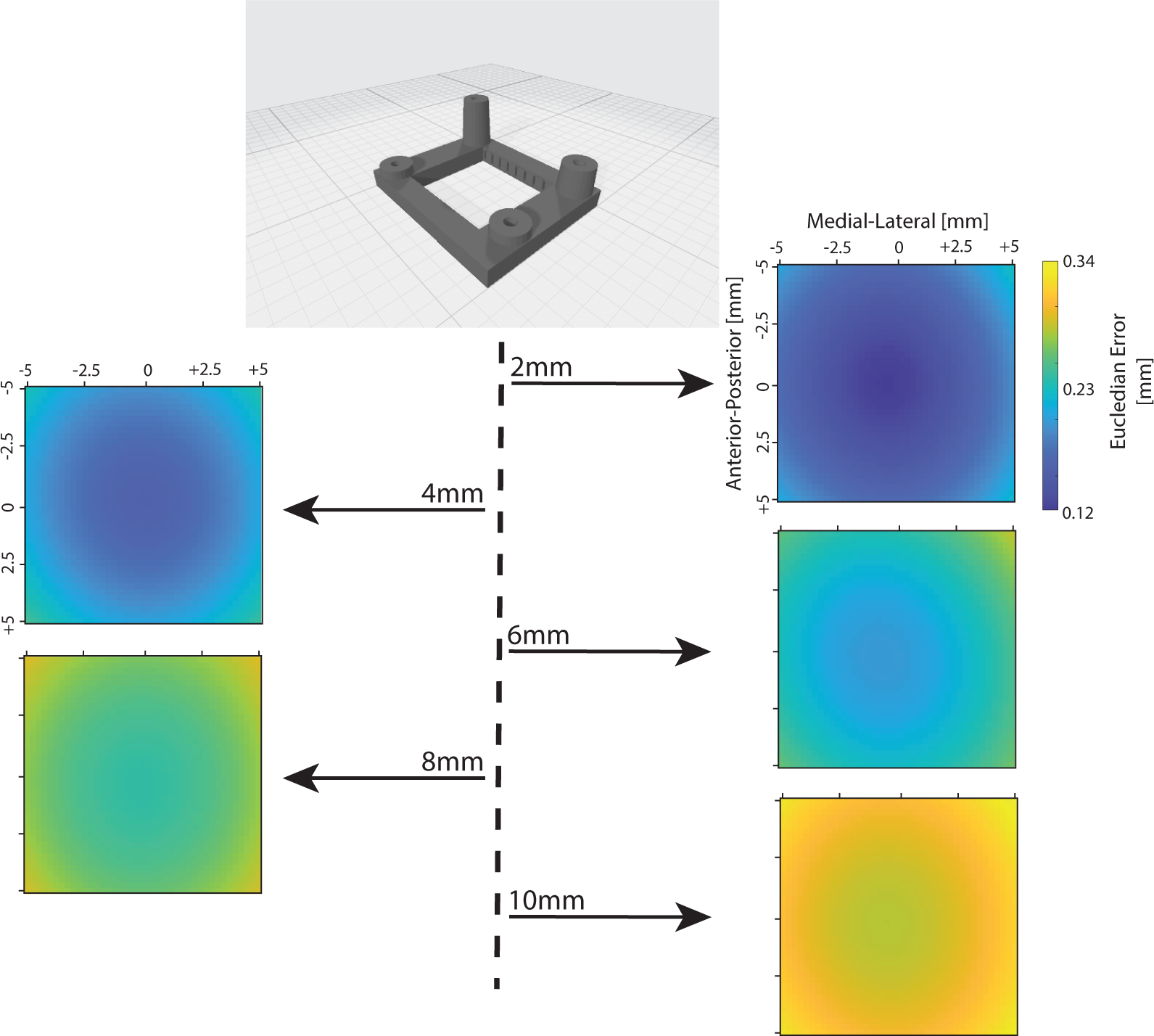
Simulation of noisy input to assess the error of target positioning. Two sets of noisy positions of fiducial markers are drawn from a normal distribution around the real positions and these are use to predict locations of targets at varying distances. The color images show the spatial distribution of the mean Eucleadian errors on 5 planes below and parallel to the marker’s frame. The mean error is color coded and it increases with distance from the center of the rigid body that is defined by the fiducial markers. At 10 mm below the frame the positioning error reaches 340 μm.

### Error assessment - physical model

To physically assess the accuracy and capabilities of the system we 3D printed a model-head at a scale similar to that of quail and mouse heads. This model incorporated the same frame for fiducial markers used in the surgeries and featured two straight ducts with different angles, each 1 mm in diameter and about 8 mm in length (Figure 6A). We then obtained a CT scan of the model-head (Figure 6B) and marked the coordinates of the four markers and the openings of the two ducts in the CT volume.

**Figure 6.**
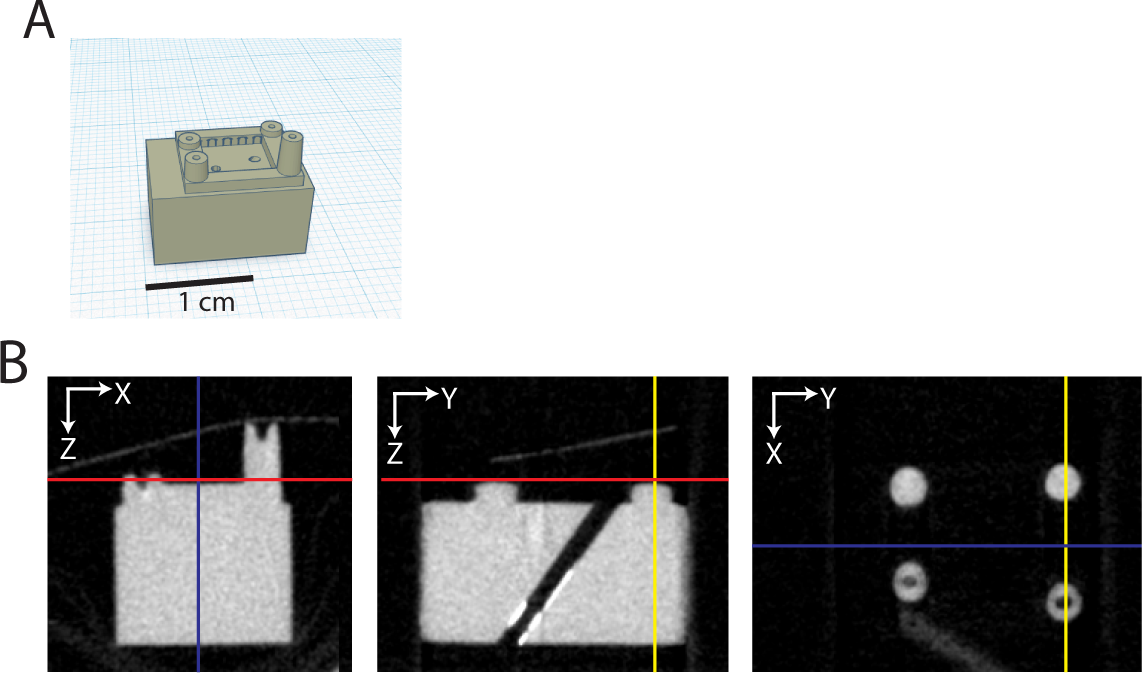
The plastic model-head used to assess the positioning capabilities of the set-up. A. A sketch of the design of the model-head. The frame used for fiducial markers in the surgery is on-top. Two ducts pass through the plastic block to serve as targets for probe penetration. B. CT images of the model-head at three orthogonal planes. In each image, the cursor lines mark the planes of the other two images.

Afterwards, the model-head was positioned in an arbitrary orientation under the XYZ positioning system. The coordinates of the fiducial markers and the openings of the two ducts were measured using the procedures described in Methods. These measurements were taken twice: once using the mechanical XYZ positioning system and once using optical tracking of the probe. For each of the 15 possible combinations of four out of six points, we calculated the transformation required to align the CT 4-points rigid body with the physical model’s 4-point rigid body, as outlined in the Methods section. We then transformed the CT coordinates of the remained two points and compared with the measured coordinates of the same points (calculating Euclidian distances between estimated and measured points).

This process was repeated at four different arbitrary orientations of the head model, resulting with 240 error estimations (15 markers combinations x 2 targets x 4 head orientations x 2 modes of measurements). The mean Euclidian error was significantly smaller when using optical tracking system to measure the fiducial markers compared to the mechanical XYZ positioning system (Figure 7A; 0.19 ± 0.09 μm and 0.24 ± 0.11 μm respectively; two sample t-test, p < 0.0001). In both measurement methods, the error along the Z-axis (dorsal-ventral) was the primary contributor to the mean Euclidean error. However, this contribution was reduced with optical tracking (Figure 8B and C; one-way ANOVA, p < 0.0001 for mechanical positioning, p = 0.016 for optical tracking).

**Figure 7.**
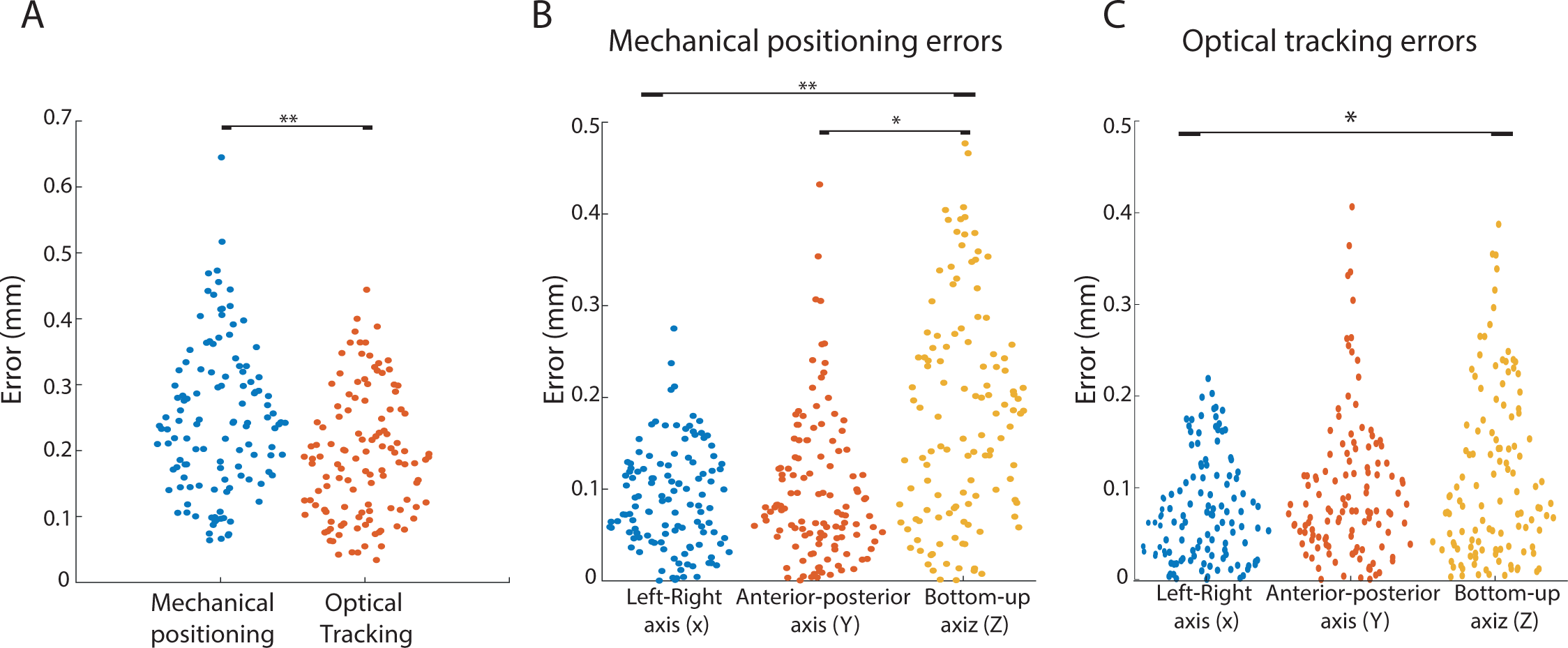
The Euclidean errors from using any 4 points in the model-head to predict the locations of the other 2 points. A. Comparison of the errors between a mechanical XYZ positioning system and an optical tracking positioning system. B. Comparison of the errors between the 3 axes of the coordinate system when using a mechanical positioning system. C. Same as in B but for the optical tracking positioning system. * designate statistical significance with a p-value smaller than 0.05; ** designate statistical significance with a p-value < 0.01 (one-tailed two samples t-test).

**Figure 8:**
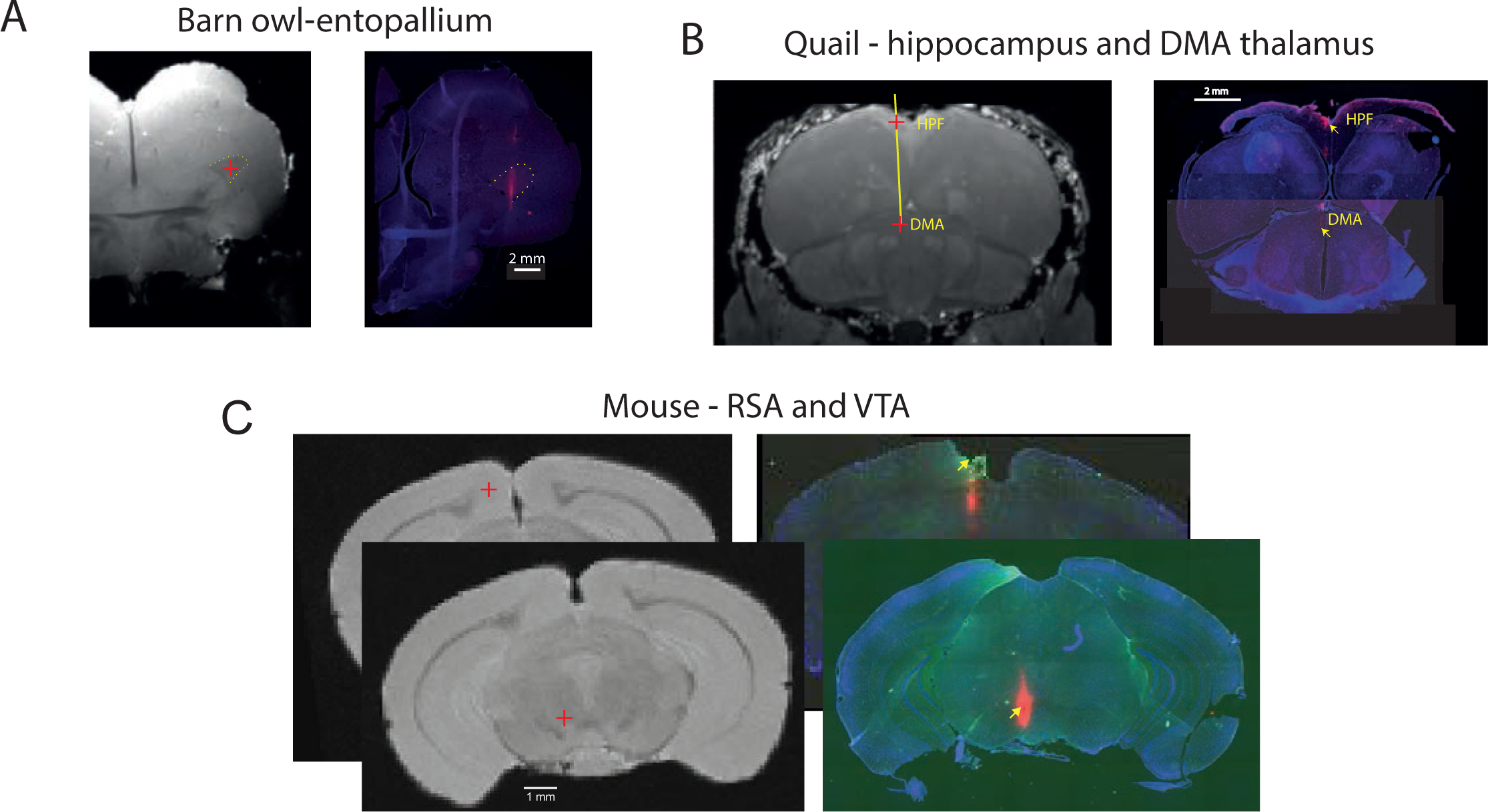
Comparison between planed and reconstructed penetration tracks. A. The picture on the left shows the MRI image of a coronal slice through the forebrain of a barn owl. The center of the entopallium is marked (red plus) as a target for the insertion of a DiI coated NPxl prob. The DAPI stained slice on the right shows that the probe was implanted in the correct brain target. B. A planed penetration track connecting two targets, the dorso-medial part of the hippocampal formation (HPF) and the dorso-medial anterior nucleus of the thalmus (DMA), is shown on the MRI slice of a quail. On the right is a Dapi stained slice of the quail’s brain. The DiI trace of the NpXl probe can be seen in the desired targets (yellow arrows). C. A penetration track connecting retrosplinial cortex (RS) with VTA of the mouse was planned (the two brain targets are marked by red pluses). The DiI traces of the NPxl can be seen in the correct targets on the right (yellow arrows).

The superior capabilities of the optical tracking system were unexpected considering the estimated error of positioning a marker through the 3D optical tracking is an order of magnitude larger than the estimated positioning accuracy of the mechanical manipulator (0.1 versus 0.01 mm respectively). However, the limiting factor here is the CT voxel size which is 0.084 mm. With optical tracking, we managed to reduce the position estimation error to below the voxel size by averaging 100 measurements of the static reflector’s position (approximately 0.06 mm, estimated from the standard deviation of repeated measurements).

To evaluate the accuracy of angle estimation and setting, we marked in the CT image the opening of the ducts at both ends as targets. Using the four fiducial markers on the frame, we calculated the angle and position of a penetration track through the two targets, starting at the upper opening and ending at the lower opening, 8 mm below. We then adjusted the angle and position of the probe and assessed the ability of the probe to clear the duct without hitting the edges. We successfully cleared the ducts with a 27G needle in different arbitrary orientations of the model-head (not quantified).

### In-vivo assessment

The ability of the system to successfully target deep brain areas for electrophysiological recordings was assessed in three animal models. First, we applied the technique in acute Neuropixels recording in the entopallium of barn owls (E). The entopallium is an apparent visual area in birds that is part of the tectofugal visual pathway (Engelage & Bischof, 1993). It is clearly distinguished in MRI and histological slices by its darker appearance from its surround (Fig 8A, dashed line), and it is located 6.5 to 8.5 mm below the brain surface. In one experiment, the Neuropixels probe was dipped in DiI (ref) and the recording track was reconstructed on bran slices (Figure 8A). The shaft clearly penetrated through the center of the E as planned in the MRI image (Figure 8A). Further verification of the success of hitting the E was obtained by recordings of single units with typical large visual RFs in the contralateral side (Reches & Gutfreund, 2009), at the expected depth of 6.5-8.5 mm below the brain surface (data not shown).

The new technique was also applied to chronically implant Neuropixels in quails. An example is shown in Figure 8B. In this example, a superficial target was defined in the hippocampal formation (HPF). The HPF lies dorsally and medial to the lateral ventricle and in the MRI is visible as lighter compared to the neural tissue below the ventricle (Fig 8B). The deeper target was defined in the dorsomedialis anterior thalamic nucleus (DMA), part of the limbic thalamus (Fig 8B, ref). The DiI trace shown in the mosaic of coronal slices in figure 8B verifies the positioning of the probe in the superficial HPF and in the DMA 4.5 mm below the HPF. Recordings in the HPF were also confirmed by the finding of a large population of head direction cells in the expected depth of the HPF (ref, data not shown).

Thus far, we successfully applied the method in two non-conventional animal models lacking established stereotaxic tools. To demonstrate the potential of using the method in the most used animal model in systems neuroscience. We used our method to implant a neuropixels dummy probe in one mouse. We aimed to connect a superficial target in the retrosplenial cortex (RSA) with a deep target in the ventral tegmental nucleus (VTA) (red Xs in Fig. 8C, left panel). The DiI trace on the DAPI stained coronal slices appears in the corresponding brain areas (yellow arrows in Fig. 8C, right panel) (ref to mouse brain atlas).

## Discussion

The traditional method for stereotaxic surgery, which involves fixing the head in a standard position within a reference frame, was developed in the early 19th century (Rahman *et al*., 2009). However, modern human brain surgery has significantly advanced, incorporating MRI imaging, fiducial markers, 3D tracking of surgical tools, and computerized tools for calculating and visualizing the surgical procedure (Kall, 1992). The advantages of these developments include surgeries tailored to an individual’s specific brain and skull, rather than relying on an average brain atlas, and freeing the head position and surgical tools from a standard, fixed frame during surgery.

In contrast, since the early days of in-vivo animal electrophysiology in the 1960s up-till today, small animal brain surgery for neuroscience research has largely adhered to the traditional approach of fixing the head in a standard position relative to a brain atlas. Nowadays, however, small animal MRI and CT scanners are increasingly accessible in research facilities (Fine *et al*., 2014), enabling the integration of techniques used in modern human brain surgery into small animal procedures. We utilized the availability of these scanners to develop a setup for stereotaxic brain surgery in small animals, inspired by contemporary human brain surgery techniques. Our method involves using a CT scan of the animal, along with fiducial markers, to align the 3D image with the actual head and brain position. This approach has been successfully employed in positioning neuropixels probes in owls, quails, and mice.

Our new method entails a more complex procedure compared to the traditional approach, owing to the additional steps of placing fiducial markers and conducting a CT scan. Consequently, it is unlikely to replace the traditional method in laboratories that use animals for which well-established stereotaxic tools exist. However, its primary advantage lies in brain research involving animals without established stereotaxic tools. For instance, barn owls have asymmetrical and intricate ear canals, rendering ear bars ineffective for keeping the head straight. Our method allows the head to be held in any convenient orientation during surgery, making it suitable for barn owls and other atypical animals.

The presented method is expected to be especially advantageous for research on animal species that have not been studied before. It facilitates the precise placement of electrodes and cannulas in the brain without necessitating extensive prior anatomical studies. A single ex-vivo MRI scan of the brain suffices.

Inspired by human brain surgery, we incorporated a 3D optical tracking device into our setup to monitor the surgical tool in real-time. While this is not essential, as fiducial markers and the probe can be positioned using a mechanical XYZ positioning system, optical tracking offers greater flexibility in probe placement. In our comparison of the accuracy in locating target positions between the mechanical and optical systems, we discovered that the optical system achieved better results. This was unexpected, given the high reliability and positioning accuracy (0.01 mm) of our mechanical positioning system (Stoelting digital mouse stereotaxic instrument). For comparison, the positioning accuracy of the 3D optical tracking system was about 0.06 mm, after averaging 100 successive measurements. We believe the superior performance of the optical tracking stems from a fundamental difference between the systems: optical tracking has a reference frame fixed to the room coordinates, whereas in mechanical positioning, the reference frame moves with the arm. This implies that the arm’s angle in the mechanical system must remain constant during the measurement of fiducial markers’ positions. Any alteration in angle will shift the coordinate system. In contrast, optical tracking permits unrestricted movement of the arm while maintaining a consistent coordinate system. This enhanced maneuverability of the probe, is likely to increase the accuracy of aligning the probe’s tip with the fiducial markers, particularly in the Z-axis, compared with the mechanical system.

More than just a specific setup, what we present is an alternative approach to small animal stereotaxic brain surgery. The setup itself can be further refined and customized to enhance precision and usability. The optical tracking device we currently use is designed for monitoring animal movements in large enclosures. Devices optimized for precise, small-volume tracking and static positioning could be more appropriate for this application, potentially offering greater affordability, ease of use, and accuracy. Additionally, exploring various manipulators, fiducial marker placements, and algorithms for optimal alignment between CT and MRI scans may yield improved results from what we have obtained.

## Acknowledgments

We thank Galit Saar and Esti Messer from the Biomedical Core Facility for their assistance and advice in MRI and CT imaging. This work was supported by research grants from the Rappaport Institute for Biomedical Research, the the Israel Science Foundation (grants no. 2655/18 and 1002/19). We also acknowledge the generous support of the Irving and Branna Sisenwine Fund.

## References

Arun, K.S., Huang, T.S. & Blostein, S.D. (1987) Least-squares fitting of two 3-D point sets. IEEE Transactions on pattern analysis and machine intelligence, 698-700.

Clark, D. & Badea, C. (2021) Advances in micro-CT imaging of small animals. Physica Medica, 88, 175–192.

De Vloo, P. & Nuttin, B. (2019) Stereotaxy in rat models: Current state of the art, proposals to improve targeting accuracy and reporting guideline. Behav Brain Res, 364, 457–463.

Dubois, A. & Bresciani, J.P. (2018) Validation of an ambient system for the measurement of gait parameters. Journal of biomechanics, 69, 175–180.

Eggert, D.W., Lorusso, A. & Fisher, R.B. (1997) Estimating 3-D rigid body transformations: a comparison of four major algorithms. Machine vision and applications, 9, 272–290.

Eilam-Altstadter, R., Las, L., Witter, M. & Ulanovsky, N. (2021) Stereotaxic brain atlas of the Egyptian fruit bat. Academic Press.

Engelage, J. & Bischof, H.J. (1993) The organization of the tectofugal pathway in birds: a comparative view. In Zeigler, H.P., Bischof, H.J. (eds) Vision, Brain, and Behavior in Birds. MIT press, Cambridge, pp. 137–158.

Fernandez, R. & Moisy, C. (2020) Fijiyama: a registration tool for 3D multimodal time-lapse imaging. Bioinformatics, 37, 1482–1484.

Fine, E.J., Herbst, L., Jelicks, L.A., Koba, W. & Theele, D. (2014) Small-animal research imaging devices. Seminars in nuclear medicine, 44, 57–65.

Güntürkün, O., Verhoye, M., De Groof, G. & Van der Linden, A. (2013) A 3-dimensional digital atlas of the ascending sensory and the descending motor systems in the pigeon brain. Brain Structure and Function, 218, 269–281.

Jun, J.J., Steinmetz, N.A., Siegle, J.H., Denman, D.J., Bauza, M., Barbarits, B., Lee, A.K., Anastassiou, C.A., Andrei, A., Aydın, Ç., Barbic, M., Blanche, T.J., Bonin, V., Couto, J., Dutta, B., Gratiy, S.L., Gutnisky, D.A., Häusser, M., Karsh, B., Ledochowitsch, P., Lopez, C.M., Mitelut, C., Musa, S., Okun, M., Pachitariu, M., Putzeys, J., Rich, P.D., Rossant, C., Sun, W.L., Svoboda, K., Carandini, M., Harris, K.D., Koch, C., O’Keefe, J. & Harris, T.D. (2017) Fully integrated silicon probes for high-density recording of neural activity. Nature, 551, 232–236.

Kall, B.A. (1992) Computer and imaging technology’s impact on stereotactic neurosurgery. A 1987–1991 update. Stereotactic and functional neurosurgery, 58, 90-93.

Karoubi, N., Segev, R. & Wullimann, M.F. (2016) The brain of the archerfish toxotes chatareus: a nissl-based neuroanatomical atlas and catecholaminergic/cholinergic systems. Frontiers in neuroanatomy, 10, 106.

Keifer, J. & Summers, C.H. (2016) Putting the “Biology” Back into “Neurobiology”: The Strength of Diversity in Animal Model Systems for Neuroscience Research. Frontiers in systems neuroscience, 10, 69.

Kleven, H., Bjerke, I.E., Clascá, F., Groenewegen, H.J., Bjaalie, J.G. & Leergaard, T.B. (2023) Waxholm Space atlas of the rat brain: A 3D atlas supporting data analysis and integration. Nature methods, 20, 1822–1829.

Lee, C.P., Xu, Z., Burke, R.P., Baucom, R., Poulose, B.K., Abramson, R.G. & Landman, B.A. (Year) Evaluation of five image registration tools for abdominal CT: pitfalls and opportunities with soft anatomy. Vol. 9413, Medical Imaging 2015: Image Processing. SPIE, City. p. 434-440.

Li, G., Xie, H., Ning, H., Citrin, D., Capala, J., Maass-Moreno, R., Guion, P., Arora, B., Coleman, N. & Camphausen, K. (2008) Accuracy of 3D volumetric image registration based on CT, MR and PET/CT phantom experiments. Journal of Applied Clinical Medical Physics, 9, 17–36.

Löffler, S., Ramrath, L., Hofmann, U.G., Schweikard, A. & Moser, A. (2009) Robot Assisted Stereotaxic Targeting for STN-DBS in the Rat Brain. Clinical Neurophysiology, 120, 49–49.

Paxinos, G., Watson, C., Pennisi, M. & Topple, A. (1985) Bregma, lambda and the interaural midpoint in stereotaxic surgery with rats of different sex, strain and weight. J Neurosci Methods, 13, 139–143.

Rahman, M., Murad, G.J.A. & Mocco, J. (2009) Early history of the stereotactic apparatus in neurosurgery. Neurosurgical Focus FOC, 27, E12.

Reches, A. & Gutfreund, Y. (2009) Auditory and multisensory responses in the tectofugal pathway of the barn owl. J Neurosci., 29, 9602–9613.

Singh, S.K., Zhang, L.B. & Zhao, J.S. (2022) Reconstruction of Flight Parameters of a Bat for Flapping Robots. Journal of biomechanical engineering, 144.

Wang, Q., Ding, S.L., Li, Y., Royall, J., Feng, D., Lesnar, P., Graddis, N., Naeemi, M., Facer, B., Ho, A., Dolbeare, T., Blanchard, B., Dee, N., Wakeman, W., Hirokawa, K.E., Szafer, A., Sunkin, S.M., Oh, S.W., Bernard, A., Phillips, J.W., Hawrylycz, M., Koch, C., Zeng, H., Harris, J.A. & Ng, L. (2020) The Allen Mouse Brain Common Coordinate Framework: A 3D Reference Atlas. Cell, 181, 936–953 e920.

Yang, R., Wang, Z., Liu, S. & Wu, X. (2012) Design of an accurate near infrared optical tracking system in surgical navigation. Journal of Lightwave Technology, 31, 223–231.

Zhou, Z., Wu, B., Duan, J., Zhang, X., Zhang, N. & Liang, Z. (2017) Optical surgical instrument tracking system based on the principle of stereo vision. Journal of biomedical optics, 22, 65005.

